# Simple mechanisms of plant reproductive benefits yield different dynamics in pollination and seed dispersal mutualisms

**DOI:** 10.1101/2021.05.05.442848

**Authors:** Kayla R. S. Hale, Daniel P. Maes, Fernanda S. Valdovinos

## Abstract

Pollination and seed dispersal mutualisms are critical for biodiversity and ecosystem services yet face mounting threats from anthropogenic perturbations that cause their populations to decline. Characterizing the dynamics of these mutualisms when populations are at low density is important to anticipate consequences of these perturbations. We developed simple population dynamic models detailed enough to distinguish different mechanisms by which plant populations benefit from animal pollination or seed dispersal. We modeled benefits as functions of foraging rate by animals on plant rewards and specified whether they affected plant seed set, germination, or negative density dependence during recruitment. We found that pollination and seed dispersal mutualisms are stable at high density but exhibit different dynamics at low density, depending on plant carrying capacity, animal foraging efficiency, and whether populations are obligate upon their partners for persistence. Under certain conditions, all mutualisms experience destabilizing thresholds in which one population declines because its partner is too rare. Plants additionally experience Allee effects when obligate upon pollinators. Finally, pollination mutualisms can exhibit bistable coexistence at low or high density when plants are facultative upon pollinators. Insights from our models can inform conservation efforts, as mutualist populations continue to decline globally.

## Introduction

Pollination and seed dispersal mutualisms support vast amounts of biodiversity and productivity in terrestrial ecosystems. Pollinators aid the reproduction of 78% and 94% of flowering plants in temperate and tropical regions, respectively (Ollerton *et al*. 2011), while seed dispersers aid the reproduction of 56% of plant species worldwide (Aslan *et al*. 2013). Unfortunately, these important mutualisms face threats globally. A recent meta-analysis of plant regeneration in forests (Neuschultz *et al*. 2016) found that pollination and seed dispersal are the processes most vulnerable to human disturbance including climate change, nutrient runoff, pesticide use, and invasive species (Stachowicz 2001, Traveset & Richardson 2006, Tylianakis *et al*. 2008, Zhou *et al*. 2013). In fact, there is clear evidence showing the abundance of pollinators and key seed disperser taxa such as frugivorous birds and mammals are declining globally (Potts *et al*. 2010, 2016, Wotton & Kelly 2011). Therefore, understanding the population dynamics of species involved in these mutualisms, especially at low abundances, can inform predictions of the consequences of those declines on the mutualisms, plant populations and crop production.

Historically, theoretical research (Gause & Witt 1935, Vandermeer & Boucher 1978, Addicott 1981, Wolin 1985, Bascompte *et al*. 2006, Okuyama & Holland 2008, Bastolla *et al*. 2009) used modified Lotka-Volterra type models (*sensu* Valdovinos 2019) to investigate the dynamics of populations interacting mutualistically. This phenomenological representation of mutualistic interactions as net positive effects between interacting species provided insight for characterizing the effects of facultative, obligate, linear, and saturating mutualisms on the long-term stability of mutualistic systems (reviewed in Hale & Valdovinos 2021). However, representing mutualisms simply as positive effects between species overlooks important dynamics that emerge from the specific mechanisms by which species positively affect each other. More recent research advanced upon these phenomenological representations of mutualism by developing models that account for consumer-resource mechanisms (Wright 1989, Holland & DeAngelis 2010, Valdovinos *et al*. 2013, Revilla 2015; see Hale & Valdovinos 2021).

Accounting for consumer-resource mechanisms enabled the discovery of important dynamics, such Allee effects (Johnson & Amarasekare 2013), alternative states (May 1976, Soberón & Martinez del Rio 1981, Wells 1983, Wright 1989, Revilla 2015), transitions between mutualism and antagonism (Holland & DeAngelis 2010), competition among species sharing mutualistic partners (Valdovinos *et al*. 2013, 2016), and niche partitioning (Valdovinos & Marsland 2021) as well as the integration of these mutualisms into food web dynamics (Hale *et al*. 2020). This research, however, mostly focused on animal dynamics (i.e., the consumer) and the dynamics of plant resources available to the animals (but see Wells 1983). To investigate the consequences of pollinator and seed disperser declines on plant populations, more focus is needed on the mechanisms by which these animals affect plant reproduction (Beckman *et al*. 2020). Here, we improve mechanistic understanding of the ecological dynamics of these mutualisms by developing and analyzing consumer-resource models that also account for simple mechanisms of reproductive benefits to plants.

Pollination and seed dispersal mutualisms share similarities as “transportation mutualisms” (Bronstein 2015). In both cases, animals visit plants to feed upon rewards (such as nectar and fruit) and provide reproductive services (such as transport of pollen and seeds) incidentally during foraging. However, these interactions differ in the mechanism of reproductive benefit to plants. Pollinators increase plant seed set by facilitating cross-fertilization, by moving pollen between conspecific plant individuals (Willmer 2011). Seed dispersers increase plant recruitment by lessening density-dependent seed(ling) mortality caused by predators, pathogens, and competitors and facilitating colonization of new habitats (Wotton & Kelly 2011, Moore & Dittel 2020). Seed dispersal can also increase germination by improving seed condition during passage through dispersers’ guts (Fricke *et al*. 2013). Our contribution investigates the dynamical consequences of these various mechanisms of reproductive benefits to plants. We find distinct dynamical behaviors at low population densities, including thresholds, Allee effects, and bistable coexistence, depending on species obligacy (i.e., their ability to persist without their partner), plant carrying capacity, and animal foraging efficiency. We also show that pollination and seed dispersal mutualisms have similar dynamics and stability at high density but differ in the ecological conditions under which the populations are vulnerable to collapse.

## Methods

We begin by deriving equations for animal mutualists, which closely follow previous consumer-resource approaches (e.g., Wright 1989, Holland & DeAngelis 2010). Then, we derive novel equations for plants, balancing simplicity with the necessary mechanistic detail to distinguish between the potential benefits of pollination versus seed dispersal interactions. We assume that the growth rate of animal and plant populations are functions of the density-independent per-capita birth (*b_A_*, *b_P_*) and death (*d_A_*, *d_P_*) rates in absence of their mutualist, where subscripts *A* and *P* indicate animal and plant populations, respectively. These population growth rates are also function of per-capita self-limitation and other density-dependent processes (*s_A_, s_P_*), and the benefits provided by mutualists (see below). Our models are continuous in time, which accommodates species with overlapping generations. They are also deterministic and ignore migration, which allows us to focus on the dynamics that emerge from the mutualistic interactions, unobscured by stochasticity and the dynamics of other patches. Table 1 summarizes our parameter definitions for all models.

**Table 1.** Table of parameters.

| Parameter | Interpretation | Units |
| --- | --- | --- |
| $a$ | Per-plant attack rate on rewards | $[P^{-1} t^{-1}]$ |
| $b_i$ | Per-capita birth or seed production rate | $[t^{-1}]$ |
| $\varepsilon$ | Efficiency of converting food rewards to animal offspring | Unitless |
| $d_i$ | Per-capita death rate | $[t^{-1}]$ |
| $f$ | Fraction of flowers pollinated in the absence of animal pollination (due to selfing, wind-pollination, etc.), $0 < f \leq 1$ | Unitless |
| $\varphi$ | Fraction of flowers pollinated due to animal-pollination, $0 < \varphi \leq 1 - f$ | Unitless |
| $g$ | Fraction of seeds that germinate in the absence of animal dispersal, $0 < g \leq 1$ | Unitless |
| $\gamma$ | Fraction of seeds that germinate due to animal dispersal (improved seed condition), $0 < \gamma \leq 1 - g$ | Unitless |
| $h$ | Handling time on rewards | $[t]$ |
| $s_i$ | Negative density-dependence due to, e.g., self-limitation during recruitment, Janzen-Connell effect | $s_P: [P^{-2}]$ or $s_A: [A^{-2} t^{-1}]$ |
| $\sigma$ | Reduced negative density-dependence due to animal dispersal (movement to safe sites), $0 < \sigma \leq s_P$ | $[P^{-2}]$ |

All parameters are assumed to be positive ( $> 0$ ). The units $A$ , $P$ , and $t$ refer to animal density, plant density, and time, respectively. Subscript $i$ is used to specify the parameter for the plant ( $P$ ) or animal ( $A$ ) population. We use Greek letters ( $\varepsilon$ , $\varphi$ , $\gamma$ , $\sigma$ ) to indicate conversion coefficients for benefits due to mutualism, while Roman letters ( $a$ , $b$ , $d$ , $f$ , $g$ , $h$ , $s$ ) indicate plant and animal demographic and consumption parameters.

### Population dynamics of animals

Pollinators and seed dispersers benefit from visiting plants by foraging for rewards that offer primarily nutritional benefit (Willmer 2011, Jordano 2014). We therefore model the change in animal population density (***A***) over time *t* as

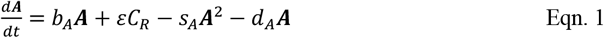

where the growth rate of the animal population increases proportionally to its consumption rate of plant rewards, *C_R_*. Parameter *ε* is the efficiency of converting rewards to new animal individuals via birth and maturation. We choose total consumption rate

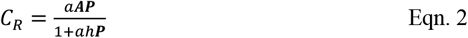

following a Holling Type II functional response with attack rate *a* and handling time *h*. This encodes the assumption that animal consumption rate on plant rewards saturates with increasing density of the plant population. Reproduction fueled by resources other than plant rewards is included in *b_A_*. We define an animal population (***A***) as an “obligate” mutualist of the plant population (***p***) when *r_A_* = *b_A_* – *d_A_* ≤ 0, that is, when ***A*** cannot persist in the absence of its mutualist plant. Otherwise (*r_A_* > 0), ***A*** is “facultative” and its population is self-sustaining.

### Population dynamics of plants

Plants benefit from reproductive services provided by animals while foraging. We define *b_P_*, the plant birth rate, specifically as maximum per-capita seed set. We assume that negative density-dependence (*s_P_*) limits the seeds that survive and mature to reproductive adults, due to, e.g., seed competition during recruitment or the preferential attraction of natural enemies (Janzen-Connell effect). Additionally, we assume that plant reproduction is limited by the fraction of flowers that can be fertilized through wind pollination or selfing (0 ≤ *f* ≤ 1) and the fraction of seeds that can germinate (0 ≤ *g* ≤ 1) even when subjected to low negative density-dependence. Reproductive services from animals increase the density of mature (reproductive) plant individuals by pollinating flowers (*φ*), improving germinability (*γ*), or providing refuge from seed predation and other sources of density-dependent mortality during recruitment (*σ*). Reproductive services are functions of animal visitation to plants, which we assume to be well approximated by the foraging rate on rewards (*C_R_*, Vázquez *et al*. 2005). Below, we derive separate models for pollination and seed dispersal based on the different mechanisms by which each affects the population dynamics of plants.

### Pollination: Increase in realized seed set from pollinating flowers

Pollination requires the transfer of pollen from one individual’s flower to the stigma of a conspecific plant individual, which occurs when animals visit flowers to forage. Visits by animals can increase the fraction of flowers that are pollinated (*f*) to a maximum of *f* + *φ* ≤ 1. We model animal-pollinated plant population dynamics (***p***) as

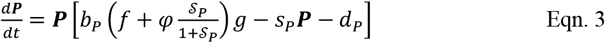

where benefit from pollination services 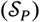 saturates as the fraction of flowers that are pollinated approaches its maximum (also see Appendix A). We assume 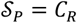, where the contribution of the animal population to per-plant seed set is the total foraging rate of animals on plant rewards (Eqn. 2). The direct dependence on plant density in this expression accounts for the repeated interactions between plant and animal individuals required for conspecific pollen transfer (Vázquez *et al*. 2005, Schupp *et al*. 2017). Using the Holling Type II functional response for *C_R_* from above and simplifying the algebra yields our specific model for animal-pollination mutualisms:

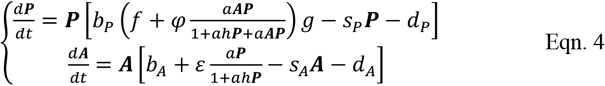

Plant benefits are limited both by the fraction of flowers available to pollinate (*φ*) and by pollinators’ handling time on flowers (*h*), with maximally effective reproductive services leading the plant population to achieve close to its maximum per-capita seed set (*b_P_*). Following our previous definitions, plant population (***P***) is an obligate mutualist of ***A*** when it cannot persist in its absence (*r_P_* = *b_P_ fg* – *d_P_* ≤ 0); otherwise, ***P*** is facultative (*r_P_* > 0).

### Seed dispersal: Reduction in negative density-dependence or increase in germination

Seed dispersers visit plants to forage on fruit, elaiosomes, or seeds, later depositing the seed away from the parent plant. This process increases recruitment by reducing density-dependent mortality caused by predators or pathogens, which are most abundant near adult plants (Wotton & Kelly 2011, Fricke *et al*. 2013, Gómez *et al*. 2019, Moore & Dittel 2020). Seed dispersal may also increase germination by improving seed condition during passage through dispersers’ guts, which removes pathogens or provides “chemical camouflage” from natural enemies (Fricke *et al*. 2013). We consider both cases of seed dispersal services 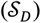 below.

First, animals rescue seeds from predators, pathogens, and intraspecific competition by transporting them away from parent plants. This can increase successful recruitment:

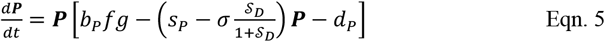

where negative density-dependence (*s_P_*) can be reduced to a minimum of *s_P_* – *σ* = 0 in the presence of animals. Eqn. 5 implies that plants must be potentially persistent (have positive demographic rates, *r_P_* = *b_P_fg* – *d_P_* >0) to benefit from reductions to negative density-dependence. Therefore, we only consider facultative plants (*r_P_* > 0) for this case.

Second, animals may increase seed condition by masticating or digesting fruits.

This can improve germinability:

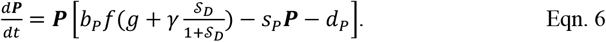

where the fraction of seeds that germinate (*g*) can increase in the presence of animals to a maximum of *g* + *γ* ≤ 1. Plants may be obligate (*r_P_* ≤ 0) or facultative (*r_P_* > 0) upon their animal partner for population persistence.

In both seed dispersal cases, reproductive services are directly related to consumption rate of animals on plant rewards. We use 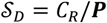, where the per-plant benefit of seed dispersal is proportional to the per-capita (i.e., per-plant) visitation rate of animals on plants. This is because repeated interactions between plant and animal individuals are possible but not required for effective seed dispersal, as they are for effective pollination (Schupp *et al*. 2017). Using the Holling Type II functional response from above yields our specific models for animal seed-dispersal mutualisms in which plants benefit via escape from negative density-dependence:

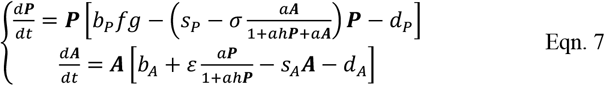

or via increased germination:

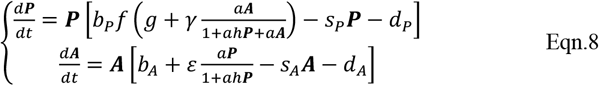

In these specific forms, plant benefits are limited both by the fraction of seeds available to germinate (*γ*) or the negative density-dependent effects to reduce (*σ*) and by dispersers’ handling time on rewards (*h*).

### Costs of mutualism

We do not model costs of mutualisms explicitly. We see three scenarios in which our choice is justified ecologically. First, individual-level costs may be already accounted for in the population-level parameters. For example, costs of producing nectar can lead to fewer ovules in flowers (Pyke 1991, Brandenburg *et al*. 2012), which reduces the ovules fertilized by animal visits and, therefore, can be accounted for by a reduced value of parameter *φ* in Eqn. 3. Second, individual-level costs may be on average (i.e., at the population-level) of such a small effect over the lifespan of the individual that they can be considered negligible. For example, nectar production is usually considered of low energetic cost (Harder & Barrett 1992, Revilla 2015). Third, other population-level parameters may drown out the effect of costs from mutualism (Ford *et al*. 2015). For example, nectar may be costly to produce for some plant species, but safe seed sites are significantly more limiting to those plants’ reproduction. Some systems may not conform to these scenarios, in which case modeling costs could be important to understand their dynamics.

### Analysis

We performed phase plane analyses on our pollination (Eqn. 4) and seed dispersal models (Eqns. 7, 8) using Wolfram Mathematica 11. Mathematical analysis and simulation code are presented in Appendix A and online, respectively. In the main text, we only present results for the ecologically relevant cases where populations have positive densities and can potentially coexist (i.e., are feasible). We investigated the behavior of populations at low density, assessing our models for conditions under which Allee effects, thresholds, and other alternative stable states occur. We use the term “Allee effects” for strong, demographic Allee effects, in which a population declines under a certain threshold of its own density (Kramer *et al*. 2009). We define “threshold effects” as population declines caused by the decline of its partner’s density under a certain threshold (Vandermeer & Boucher 1978, Revilla 2015). Alternative stable states, such as single-species persistence or bistable coexistence, occur when a system can settle stably into more than one equilibrium depending only upon initial conditions (“historical accidents,” May 1977).

## Results

We found that pollination and seed dispersal mutualisms have similar dynamics and stability at high density but differ in the ecological conditions under which the populations are vulnerable to collapse. Different parameter regimes, in combination with whether each partner is obligate or facultative, lead to different dynamics at low population densities. Below, we describe these different dynamics, assuming that coexistence is feasible (but see Appendix A for mathematical details). Then, we describe ecological scenarios under which each outcome is likely to occur. We provide these scenarios rather than formal mathematical conditions, because the latter are difficult to interpret due to the nonlinearities in our models.

### Pollination

The shape and intersections of the plant and animal “nullclines” (curves of zero change in population density) determine the dynamics of the system. Both species have “trivial” nullclines at zero density. The non-trivial animal nullclines (black curves) are concave down, increasing functions that saturate with respect to plant density (Figs. 1-3). The non-trivial animal-pollinated plant nullclines (green curves) are U-shaped when plants are obligate mutualists (Fig. 1A-B) or increasing and concave up at high densities when plants are facultative (Fig. 1C-G). Depending on the parameter regime, the facultative plant nullcline may exhibit an inflection point at low densities in which it transitions from concave down to up (e.g., Fig. 1D). Facultative animals and plants may persist in the absence of their partners at density *K_A_* = *r_A_/s_A_* and *K_P_* = *r_P_/s_P_*, respectively. These densities are “carrying capacities” in the original sense of single-species equilibrium densities (Vandermeer & Boucher 1978). When coexistence occurs, it is at a higher density than either species could achieve alone (> *K_P_*, > *K_A_*).

**Figure 1.**
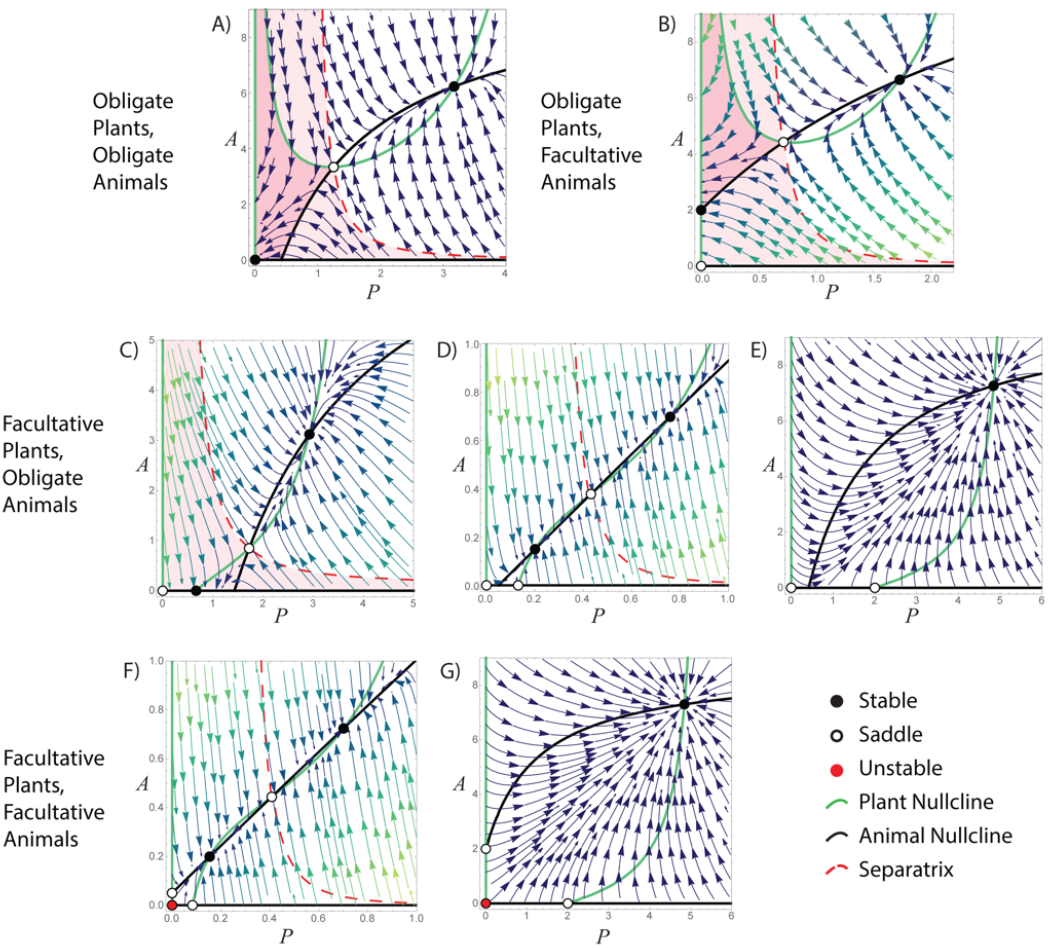
Phase plane diagrams for plant-pollinator mutualisms. Nullclines are curves of zero growth for plant (green) or animal pollinator (black) populations. Equilibria (dots) occur at the intersection of plant and animal nullclines. Filled black equilibria are stable attractors, filled red equilibria are unstable repellers, and hollow equilibria are saddle points, which repel in one dimension and attract in the other. Arrows show the directions of population change for plants (x-axis) and animals (y-axis), with lighter colors indicating a faster rate of change. Dynamics depend upon the parameterization of the model and whether plants and animals are obligate or facultative mutualists (rows). When plants are obligate (**A, B**), they experience Allee effects at low density (dark red shaded region). Plants cannot attract sufficient pollinator visitation to support their own population growth, leading to collapse. When either plants or animals are obligate, both partners experience threshold effects (light red shaded region). Below the threshold marked by the separatrix (dashed line), one population’s density is too low to support its partner’s growth, resulting in system collapse to extinction (**A**), animal-only persistence at *K_A_* = (*b_A_* – *d_A_/s_A_* (**B**), or plant-only persistence at *K_P_* = (*b_P_fg* – *d_P_*)/*s_p_* (**D**). Thus, depending upon the initial densities of both partners, the mutualism may persist stably or collapse. When one partner is facultative, stable coexistence may be the only outcome (**C, E, G**). However, in a small parameter space, bistable coexistence may occur (**D, F**). Here, the separatrix divides the region in which the system will be attracted to the low- or high-density stable coexistence equilibrium, depending upon whether initial densities are below or above the separatrix, respectively. Parameter values are fixed to the following: *b_P_* = 1, *f* = 1, *φ* = 0.5, *g* = 1, *s_p_* = 0.15, *a* = 0.8, *h* = 1, *b_A_* = 1, *ε* = 2, *s_A_* = 0.15, except for: (**A**) *s_P_* = 0.05, *d_P_* = 0.75, *d_A_* = 1.5; (**B**) *b_P_* = 1.2, *d_P_* = 0.76, *a* = 0.31, *d_A_* = 0.7; (**C**) *s_P_* = 0.14, *d_P_* = 0.4, *a* = 0.9, *s_A_* = 0.115, *d_A_* = 2.1; (**D**) *b_P_* = 1.5, *s_P_* 0.61, *d_P_* = 0.67, *a* = 2, *h* = 0.01, *ε* = 1, *s_A_* = 2, *d_A_* = 1.1; (**E**) *d_P_* = 0.2, *d_A_* = 1.5; (**F**) *b_P_* = 1.5, *s_P_* = 0.6, *d_P_* = 0.7, *a* = 1.95, *h* = 0.01, *ε* = 1, *s_A_* = 2, *d_A_* = 0.9; (**G**) *d_P_* = 0.2, *ε* = 1, *d_A_* = 0.7.

**Figure 2.**
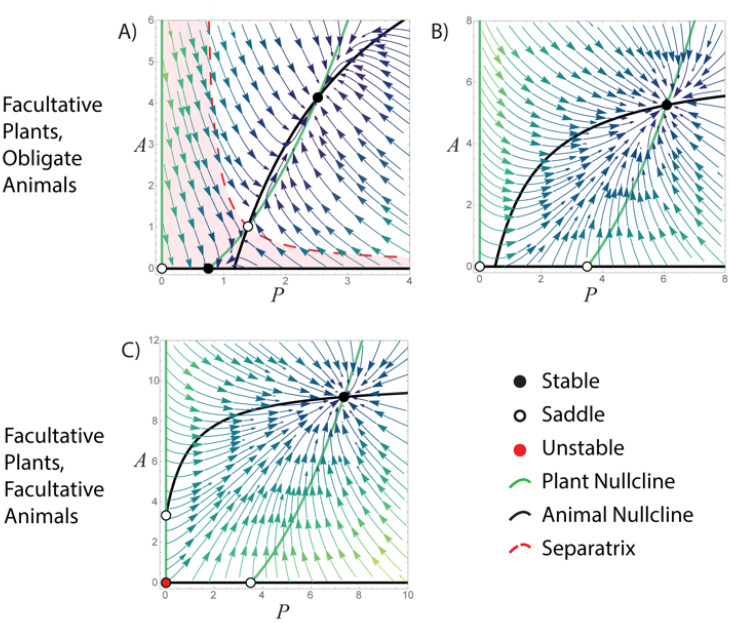
Phase plane diagrams for plant seed-disperser mutualisms, where animals reduce negative density-dependence in plants. Nullclines are curves of zero growth for plant (green) and animal seed-disperser (black) populations. Here, plants must be able to persist at low density (*K_P_* > 0) to benefit from reductions in negative density-dependence; therefore, we consider only facultative plant populations. All other formatting and terminology follow Fig. 1. When animals are obligate mutualists (**A**), both partners may experience threshold effects. Otherwise, both partners experience growth from low density (**B, C**), resulting in a single stable coexistence equilibrium. Parameter values are fixed to the following *b_P_* = 1, *f* = 1, *g* = 1, *s_P_* = 0.2, *σ* = 0.2, *d_P_* = 0.3, *a* = 1, *h* = 1, *b_A_* = 1, *s_A_* = 0.15, except for: (**A**) *d_P_* = 0.85, *a* = 2, *h* = 0.5, *b_A_* = 1, *ε* = 0.93, *s_A_* = 0.08, *d_A_* = 2; (**B**) *ε* = 1.5, *d_A_* = 1.5; (**C**) *ε* = 1, *d_A_* = 0.5.

**Figure 3.**
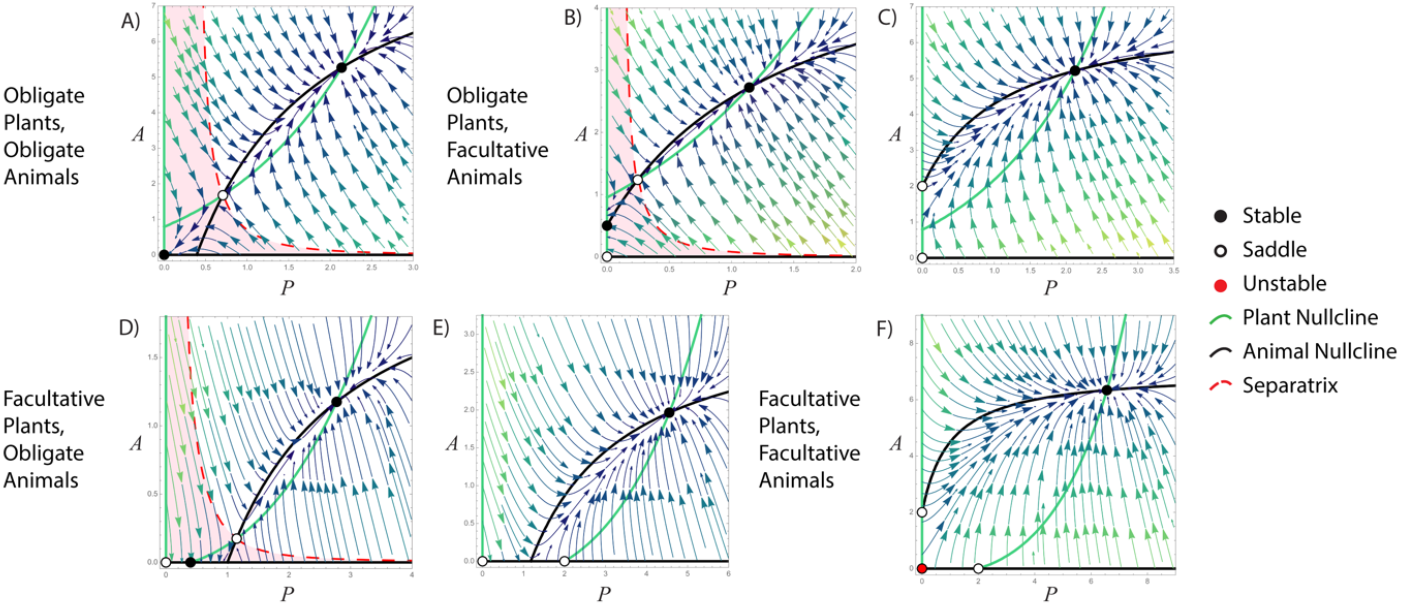
Phase plane diagrams for plant seed-disperser mutualisms, where animals increase seed germination in plants. Formatting and terminology follow Fig. 1. When either partner is an obligate mutualist, both partners may experience threshold effects (**A, B, D**). When either partner is a facultative mutualist, the system may instead exhibit a single coexistence equilibrium with growth from low density for both partners (**C, E, F**). Parameter values are fixed to the following *b_P_* = 1, *f* = 1, *g* = 0.5, *γ* = 0.5, *s_P_* = 0.05, *d_P_* = 0.7, *a* = 0.85, *h* = 1, *b_A_* = 1, *ε* = 1, *s_A_* = 0.2, *d_A_* = 1.5, except for: (**A**) *ε* = 2, *s_A_* = 0.15; (**B**) *s_A_* = 0.7, *d_A_* = 0.9; (**C**) *d_A_* = 0.6; (**D**) *d_P_* = 0.48, *a* = 1; (**E**) *d_P_* = 0.4, *s_A_* = 0.15; (**F**) *a* = 1, *d_A_* = 0.6.

When plants are obligate (Fig. 1A-B), pollination mutualisms exhibit two non-trivial equilibria: First, a stable coexistence equilibrium (off-axes filled circle). Density past this equilibrium in either species causes its population to decrease due to negative density-dependence. Second, a saddle point that attracts in one dimension but repels in the other (off-axes hollow circle), bisecting the plane into two portions, as marked by the “separatrix” (dashed line). This separatrix marks out a threshold under which obligate partners go extinct even if initially highly abundant (“threshold effects,” regions shaded light red). For example, following a trajectory from the lower-right region in Fig. 1B, initially highly abundant obligate plants go extinct while initially rare pollinators persist at *K_A_*. These threshold effects occur because one species is too low in density to provide sufficient benefits to its partner, causing the partner’s population to decline. The low-density species continues to benefit from mutualism but its increase in density cannot occur fast enough to save the system from collapse. Above the separatrix, one or both species are of high enough density that benefits from mutualism cause positive population growth in their partners and the system will achieve stable coexistence.

Obligate animal-pollinated plants are additionally susceptible to Allee effects (regions shaded dark red), where their population declines under a threshold of their own density regardless of the density of their partner (Fig. 1A-B). This occurs because benefit is proportional to the total consumption rate by animals, i.e., plants require obligate outcrossing for successful pollination behavior. At very low plant density, even highly abundant pollinators are limited by the time required to harvest and (incidentally) transfer pollen between two plant individuals, and thus cannot provide pollination services at a rate sufficient to allow plant population growth. Unsurprisingly, animals cannot acquire enough food and decline either to extinction (Fig. 1A) or to *K_A_* (Fig. 1B) when plants are extinct.

When animal-pollinated plants are facultative (Fig. 1C-G), more diverse dynamics are possible. High-density stable coexistence always occurs if partners are originally at high enough density. At lower density, threshold effects can occur when animals are obligate mutualists (Fig.1C), but population growth from low densities is also possible (Fig. 1D-G) and even guaranteed when both partners are facultative mutualists (Fig. 1F-G). Additionally, in a small parameter range, the mutualism can exhibit bistable coexistence (Fig. 1D, 1F). Here, there are two possible points of stable coexistence which the system will be attracted to – one at lower density and one at higher – based on initial conditions. A separatrix (dashed line) running through a central saddle point (hollow circle) marks out a threshold under which the system will persist at the lower-density coexistence equilibrium.

By inspecting the intercepts of the nullclines with the axes, we gain intuition into ecological scenarios when these different dynamics (threshold effects, stable coexistence, or bistable coexistence) will occur. Threshold effects (Fig. 1C) can occur when 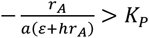, that is, when the intercept of the animal nullcline with the x-axis (representing plant density) is greater than the intercept of the plant nullcline. This is most likely when pollinators are highly obligate upon plants (*r_A_* ≪ 0) and when plants have low carrying capacity, *K_P_*. That is, when the environment is hostile due to, for example, high density-independent mortality (*d_P_*, *d_A_*) or high negative density-dependence for plants (*s_P_*). On the other hand, growth from low density (Fig. 1D-G) can occur under the opposite condition, when 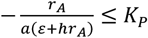. That is, when the intercept of the animal nullcline with the x-axis is less than the intercept of the plant nullcline. This is most likely when animals are facultative or only weakly obligate and plants have high *K_P_*. However, if this condition holds but plants have low *K_P_*, bistable coexistence may occur. Bistable coexistence is most likely when pollinators have near-zero population growth rate (*r_A_*) in the absence of plants, regardless of whether they are obligate or facultative.

The potential for bistable coexistence is strongly modulated by pollinators’ foraging efficiency, especially their attack rate (*a*) or search time (1/*a*) for floral rewards (Fig. 4A). At higher *a*, only a single, high-density, coexistence equilibrium is present. At lower *a*, stable coexistence occurs only at low density or is infeasible. However, at intermediate *a*, bistable coexistence can occur. Pollinators can find plants but do so inefficiently enough that the plant and animal populations never achieve a high enough growth rate to escape a low-density stable attractor, where plants stay relatively rare, and animals stay inefficient at finding them. This happens unless initial plant abundance is high enough to be easily found by animals so that sufficient mutualistic benefits can be exchanged to allow stable coexistence at high density.

**Figure 4.**
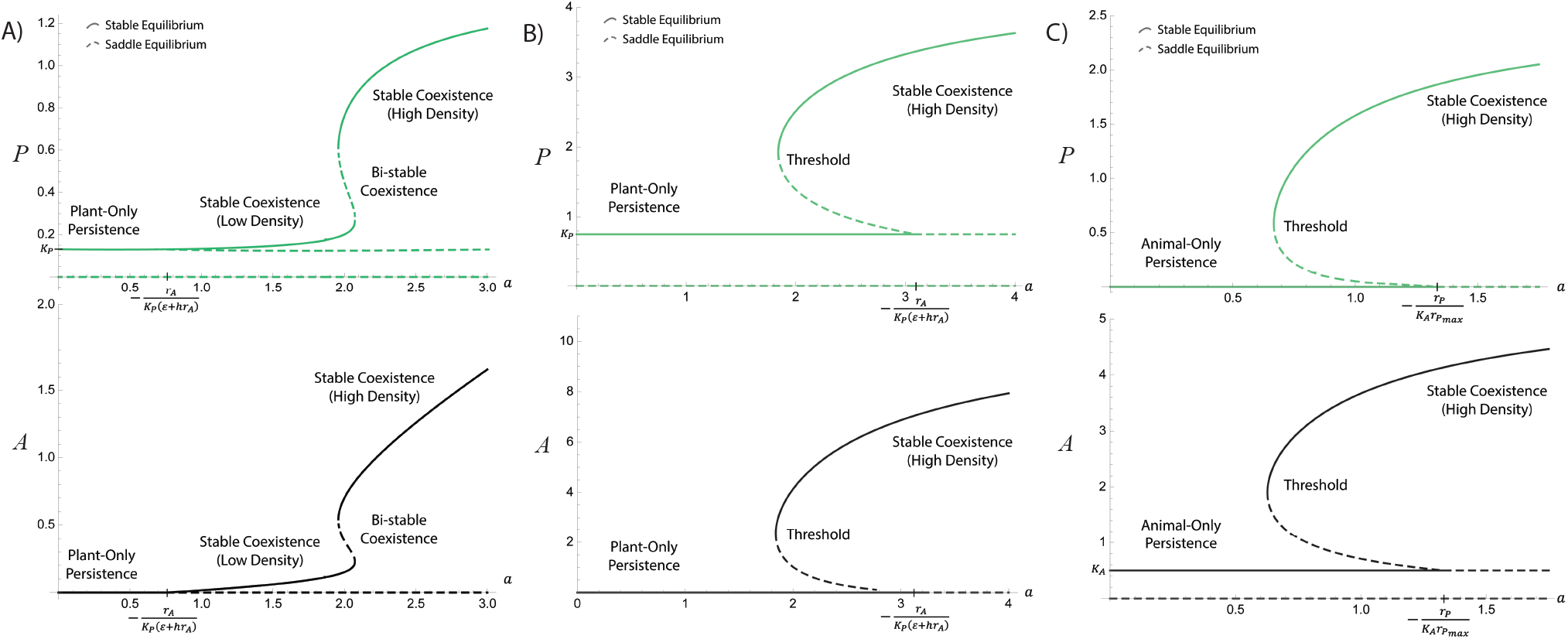
Bifurcation diagrams: animal foraging efficiency (*a*) mediates the dynamics of pollination and seed dispersal mutualisms. Green (top) and black (bottom) curves show plant and animal equilibrium density, respectively, across a range of animal foraging efficiencies or attack rate on plants’ rewards (*a*, x-axis). This parameter mediates both animal benefits from plants (foraging on rewards) and plant benefits from animals (foraging rate determines the rate at which plant gametes are transported). When multiple curves occur at the same *a* value, multiple equilibria are possible. Assuming feasible coexistence: (**A**) Pollination mutualisms with facultative plants: At low *a*, stable coexistence occurs at low density for both partners, whereas at high a, stable coexistence occurs at high density. At intermediate *a*, coexistence occurs at either low or high density (bistable coexistence) depending on the initial densities of the mutualists (whether densities are below or above the separatrix in Fig. 1D, 1F). (**B**) Seed dispersal with obligate animal mutualists that reduce plant negative density-dependence: At low *a*, stable coexistence occurs if populations are initially at high density. Below a threshold in initial density (separatrix in Fig. 2A), the mutualism collapses and animals go extinct. As *a* increases, this threshold becomes lower and lower until populations beginning at any positive density exceed it (when 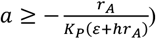, ensuring stable coexistence. Pollination mutualisms and seed dispersal mutualisms that increase germination also exhibit these dynamics. (**C**) Seed dispersal mutualisms that increase germination of obligate plants: The transition from threshold effects to stable coexistence occurs when 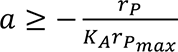. Saddle point equilibria are dashed lines, while stable equilibria are solid. In all cases, extinction of both partners and single-species persistence of facultative partners (at *K_i_*) are possible equilibria. Coexistence is infeasible at very low *a*, permitting only single-species persistence or extinction. Note that the inverse of animal attack rate on plants’ rewards (1/*a*) can be interpreted as animals’ search time for plant rewards, where higher search time could be caused by limitations of the animals’ sensory ability for identifying available food on the landscape. Attack rate can also be interpreted as animals’ preference, where lower preference could result from lower quality of the provided rewards or risk associated with accessing them. Parameters are fixed to: (**A**) *b_P_* = 1.5, *f* = 0.5, *φ* = 0.5, *g* = 1, *s_P_* = 0.61, *d_P_* = 0.67, *h* = 0.01, *b_A_* = 1, *ε* = 1, *s_A_* = 0.2, *d_A_* = 1.1; (**B**) *b_P_* = 1, *f* = 1, *g* = 1, *s_P_* = 0.2, *σ* = 0.2, *d_P_* = 0.85, *h* = 0.5, *b_A_* = 1, *ε* = 0.93, *s_A_* = 0.08, *d_A_* = 2; (**C**) *b_P_* = 1, *f* = 1, *g* = 0.5, *γ* = 0.5, *s_P_* = 0.05, *d_P_* = 0.7, *h* = 1, *b_A_* = 1, *ε* = 1, *s_A_* = 0.2, *d_A_* = 0.9, while *a* is varied.

### Seed Dispersal

Our seed dispersal models (Figs. 2-3) display simpler dynamics than our pollination model. Seed dispersers follow the same dynamics as pollinators (black curves). The non-trivial nullclines for seed-dispersed plants are concave up, increasing functions (green curves), bounded by a vertical asymptote on the right. This results in stable coexistence at higher density than either species could achieve alone with the potential for threshold effects, but not the bistable coexistence or Allee effects dynamics described for pollination mutualisms.

When seed dispersers benefit plants through reduced negative density-dependence (Fig. 2), plants are always facultative (see Methods) and can persist in the absence of animal mutualists at density *K_P_*. The same qualitative conditions hold as described for the pollination model as to whether the system exhibits threshold effects 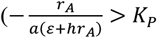, Fig. 2A) or a single stable coexistence equilibrium 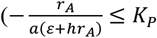, Fig. 2B) when animals are obligate mutualists.

When seed dispersers benefit plants through increased seed germination (Fig. 3), plants may be obligate or facultative, with population collapse or persistence at *K_P_*, respectively, in the absence of animals. When at least one partner is an obligate mutualist, threshold effects can always occur, leading to extinction if at least one partner is at low enough density (Fig. 3A-B, 3D). As shown for the previous models, threshold effects can occur when 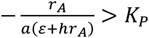 (Fig. 3A, 3D). They can also occur in this model when 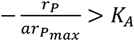, where *r_P_max__* = *r_P_* + *b_P_fγ*. That is, when the intercept of the plant nullcline with the y-axis (representing animal density) is greater than the intercept of the animal nullcline (Fig. 3A-B).

In both seed dispersal cases, threshold effects are most likely when one partner is highly obligate (*r_i_* ≪ 0) and when plants are difficult, but not impossible, for dispersers to find on the landscape (low search time, 1/*a*, Fig. 4B-C). At even lower *a*, coexistence is infeasible while at higher a, dispersers are adept at finding plants, and both populations grow quickly to high density stable coexistence. High density stable coexistence is also possible if neither partner is highly obligate and dispersers are efficient at finding plants (Fig. 3C, 3E). This is the only outcome when both partners are facultative (Fig. 2C, 3F).

## Discussion

Pollination and seed dispersal mutualisms support the reproduction of a vast number of plant species globally (Aslan *et al*. 2013, Neuschultz *et al*. 2016). Anthropogenic stressors threaten these mutualisms by causing population declines that can disrupt the benefits provided by the mutualists (Tylianakis *et al*. 2008, Traveset & Richardson 2006, 2014, Potts *et al*. 2010, 2016, Beckman *et al*. 2020). These potential disruptions increase the need to study the dynamics of mutualisms when the interacting populations exhibit low density. Theoretical studies have traditionally investigated these dynamics at low density by using Lotka-Volterra models. Our work builds from those studies by developing novel consumer-resource models of mutualistic interactions (*sensu* Holland & DeAngelis 2010) that incorporate simple mechanisms of reproductive benefits to plant populations. Our results expand the understanding of pollination and seed dispersal mutualisms by elucidating dynamical consequences of mechanisms by which pollinators and seed dispersers benefit plant reproduction when visiting them as consumers.

We found that both pollination and seed dispersal mutualisms exhibit threshold effects when at least one mutualist is obligate (Fig. 5). Threshold effects, population declines due to rarity of a mutualistic partner, were first described for obligate mutualists that strongly benefit each other in studies using the Lotka-Volterra model (May 1976, Vandermeer & Boucher 1978). More recent models incorporating consumer-resource mechanisms also predict these thresholds (May 1976, Soberón & Martinez del Rio 1981, Wells 1983, Wright 1989, Fishman & Hadany 2010, Revilla 2015). However, despite their ubiquity in theoretical work (reviewed in Hale & Valdovinos 2021), threshold effects have been difficult to observe empirically (Latty & Dakos 2019, Hillebrand *et al*. 2020). Our results may explain this difficulty by suggesting that threshold effects would only be observed in pollination systems inhabiting hostile environments (low carrying capacity, Figs. 1C, 5) when pollinators are obligate and plants are facultative. Otherwise, destabilization of the system would be attributed to Allee effects in plants (Fig. 5). Our results also show that threshold effects may be easier to observe in seed dispersal systems because they would manifest when either partner’s density drops below the critical threshold. As a potential example, Wotton and Kelly (2011) observed that seed survival of two species of New Zealand trees dropped dramatically when fruit consumption by dispersers crossed below 30%.

**Figure 5.**
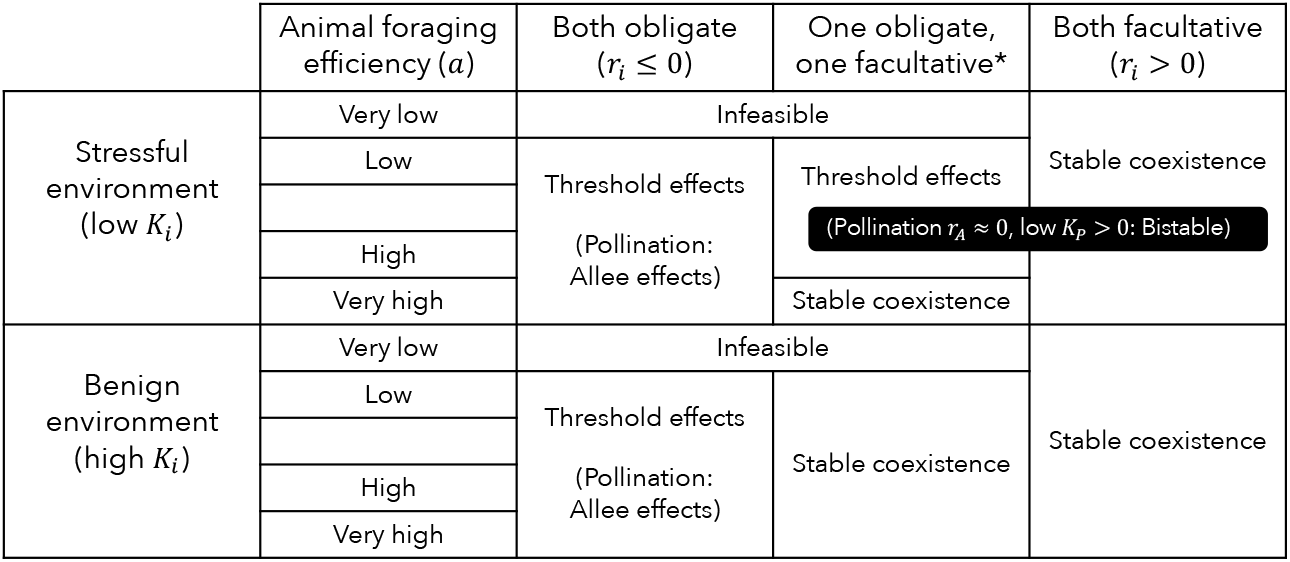
Summary diagram of our results: pollination and seed dispersal dynamics under different ecological scenarios. Critical transitions between different dynamics are strongly controlled by both partners’ obligacy (governed by per-capita growth rate, *r_i_*), the level of environmental hostility (governed by carrying capacity, *K_i_* = *r_i_*/*s_i_*), and animals’ foraging efficiency on rewards (or attack rate, *a*). Species may be facultative mutualists (*r_i_* > 0) or obligate upon their partners for persistence (*r_i_* ≤ 0). In hostile environments, facultative and obligate mutualists are assumed to exhibit low and very negative “carrying capacities,” respectively. In benign environments, facultative or obligate species have high or barely negative carrying capacities, respectively. At very low animal foraging efficiency, coexistence is infeasible if at least one species is obligate. When both species are facultative, stable coexistence always occurs when feasible, regardless of obligacy or environmental hostility. When both species are obligate, threshold effects (and Allee effects for pollination mutualisms) occur when coexistence is feasible. In seed dispersal mutualisms: i) threshold effects occur in hostile environments when at least one species is obligate, unless attack rate is very high in which case stable coexistence may occur, and ii) stable coexistence always occurs in benign environments when at least one partner is facultative. The asterisk indicates that in pollination mutualisms, i) and ii) occur specifically when plants are the facultative partner and animals are obligate. Otherwise, threshold effects and Allee effects occur when animal-pollinated plants are obligate, regardless of animals’ obligacy and the environmental condition. Pollination mutualisms uniquely exhibit bistability when plants are facultative and *r_A_* is close to zero, regardless of animal obligacy.

Allee effects may also cause irrecoverable declines in populations at low density. We use “Allee effects” to describe self-induced, as opposed to partner-induced, declines at low density. We find Allee effects in plant populations that are obligate mutualists of pollinators, due to, e.g., high self-incompatibility. This result is consistent with pollination ecology because pollinators’ benefits to plants tend to decrease at low plant density due to increased search time, leading to a decrease in visitation rate and pollen limitation (Forsyth 2003). Low plant density may also decrease visitation rate due to insufficient floral displays or reduce visit quality due to pollen dilution (Forsyth 2003). Indeed, pollinator-mediated Allee effects have been observed empirically in highly self-incompatible plant species (Kramer *et al*. 2009). We show that reductions in visit quantity alone are sufficient to induce an Allee effect, whereas natural populations likely experience both a decrease in visit quality and quantity at low plant density (Kunin 1993). In contrast, we did not find Allee effects in our seed dispersal models. This is consistent with the mechanisms by which seed dispersers benefit plants. Seed dispersers benefit plants by increasing the seed germination or their survival to adulthood through refuge from intraspecific competition and natural enemies (seed predators, herbivores) that are attracted to regions of high food densities (i.e., the Janzen-Connell effect, Janzen 1970, Connell 1971, Wotton & Kelly 2011, Fricke *et al*. 2013, Moore & Dittel 2020). These plants are thus less likely to experience Allee effects, because at low density the primary threats are similarly reduced.

We found bistable coexistence for our pollination models, where partners coexist stably either at low or high density, depending upon if the populations dip below a certain threshold of density, especially driven by plant density. Specifically, for the case of facultative plants inhabiting a hostile environment (low carrying capacity) whose pollinators are of intermediate foraging efficiency. It is, to our knowledge, a novel result in mutualism models (but see Zhang 2003 and Holland & DeAngelis 2010 for examples when mutualism can transition dynamically to competition or parasitism interactions). We are not aware of direct, empirical evidence for bistable coexistence. However, we propose that the mechanism we found here for bistability to occur (i.e., benefits plants accrue from pollinator visits increase with plant density) may be inferred from previous empirical observation. In particular, from the observation that plant abundance explains the variability of total benefits that populations of facultative plants receive from pollinator visits (Vázquez *et al*. 2007).

Environmental hostility or stress has received much attention as a potential cause for tipping points (Beisner *et al*. 2003, Lever *et al*. 2014, Kéfi *et al*. 2016, Latty & Dakos 2019, Hillebrand *et al*. 2020, Huang & D’Odorico 2020), which we also find here (Fig. 5). We find that species can transition from “facultative” to “obligate” mutualists with increasing death rate, potentially leading to low-density thresholds. Additionally, lower carrying capacity may cause a species to exhibit only low-density or infeasible coexistence. We also find that animal foraging efficiency affects critical transitions between these dynamics (Fig. 5). In particular, decreased foraging efficiency can cause a previously stable system at high density to shift to bistability, threshold effects, or collapse to infeasible coexistence. Decreased foraging efficiency can occur with increased crypticity of rewards due to habitat fragmentation, poor attack rate due to mismatched traits, or low preference due to declines in rewards quality. These results are consistent with previous work by Valdovinos and Marsland (2021) who identified a threshold for quality of visits below which the plant species receiving those visits and the animals depending on those plants go extinct. This suggests that the parameter of foraging efficiency (analogous to visit quality in their model) could be an important topic for future investigation.

A unique aspect of our modelling approach is that we specify whether plants benefit according to animal total or per-capita visitation rate. Our choice for pollination benefits to plants to scale with animal total visitation rate accounts for outcrossing (Vázquez *et al*. 2005). This, to the best of our knowledge, makes our pollination model unique in the literature of two-species models (but see Valdovinos *et al*. 2013, 2016, Hale *et al*. 2020 for network models). In addition, visitation in our models differs from previous work (e.g., Revilla 2015) because we use nonlinear (specifically, Holling Type II) saturating functional responses for consumption rate. This allows both plants and animals to experience intra-specific competition for benefits, set by the maximum time animals can spend foraging on rewards. Lastly, we impose a direct limitation to plant benefits, so that they saturate due to limited ovules or seeds. In this way, plant benefits in our seed dispersal models follow the Beddington-DeAngelis functional response, which was formulated for consumers experiencing exploitation competition.

Our models can also accommodate functional responses interpreted as “net benefit” curves (e.g., Holland *et al*. 2002, Morris *et al*. 2010) if costs and benefits affect the same vital rate. That is, if “costs” simply reduce the benefits that accrue to a given vital rate (e.g., Brandenburg *et al*. 2012). Under this interpretation, unimodal functional forms may arise (e.g., Morris *et al*. 2010) which could lead to substantially different dynamical predictions than those presented here. Finally, our models, as well as previous ones, assume that mutualisms have population-level impacts. Most empirical studies, however, quantify the benefits and costs of mutualisms at the individual level in terms of fitness or even by using a single proxy for fitness (Bronstein 2001, Ford *et al*. 2015). Those effects do not necessarily imply population- or community-level impacts of mutualism (Flatt & Weisser 2000, Ford *et al*. 2015). Therefore, empirical work on population dynamics of mutualisms is of foremost importance to evaluate historical and current ecological theory on mutualisms.

## Conclusion

This research increased mechanistic understanding of the dynamics of plant-animal mutualisms by analyzing consumer-resource models that incorporate simple mechanisms of reproductive benefits to plants. We found that these mutualisms may be vulnerable to declines at low density due to Allee and threshold effects and identified potential bistable coexistence. We also characterized the ecological scenarios under which those dynamical behaviors are most likely to occur, which has been recognized a conservation priority (Latty & Dakos 2019). Future work should continue to investigate how different mechanisms of mutualism manifest in different population dynamics, both to advance ecological understanding and to aid in conservation objectives.

## Acknowledgements

This research was supported by National Science Foundation Graduate Research Fellowship DGE-1143953 to K.R.S.H., National Science Foundation Graduate Research Fellowship DGE-1256260 to D.P.M., and National Science Foundation grant DEB-1834497 to F.S.V.

## Statement of authorship

F.S.V. conceived the study; K.R.S.H. and D.P.M. analyzed the models; K.R.S.H. and F.S.V. developed the biological interpretation of the model results; K.R.S.H. wrote the manuscript with important contributions from F.S.V.; all authors developed the models.

## Code availability

Mathematica notebooks used to analyze the models are available at https://doi.org/10.5281/zenodo.5787056

## Appendix A: Mathematical Analysis

In all our models (Eqns. 4, 7, 8), plant and animal populations have “trivial” nullclines: 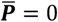, a vertical line along the y-axis, and 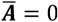, a horizontal line along the x-axis. The intersection of these trivial nullclines results in a trivial extinction equilibrium 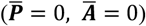 in each model. Below, we describe the geometry of the nontrival nullclines (hereafter, simply “nullclines”) in the ecologically relevant region of the plane, when ***P*** ≥ 0, ***P*** ≥ 0 (hereafter, simply “the positive quadrant”). All our models are structurally unstable (Rohr *et al*. 2014), such that smooth transitions in parameter values shift the nullclines so that they may intersect in various ways or even fail to intersect in the positive quadrant, with different dynamical outcomes for each case (Fig. 4). Below, we also describe how the nullcline geometries are dependent on obligacy to mutualism, which leads to structural instability.

The animal population follows the same dynamics (Eqn. 1) and thus has the same nontrivial nullcline in all our models (Figs. 1-3, black):

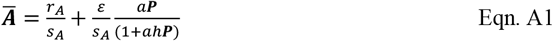

where *r_A_* = *b_A_* – *d_A_*. This equation describes the balance between negative density-dependence, benefit from mutualism, and intrinsic growth or decay for facultative (*r_A_* > 0) or obligate (*r_A_* ≤ 0) animal populations, respectively. When 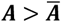, negative density-dependence is stronger than benefit from mutualism and/or intrinsic growth, causing ***A*** to decrease. When 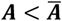, negative density-dependence is weaker and ***A*** increases.

Specifically, Eqn. A1 is a concave down, increasing function that saturates with respect to plant density, ***p***. The benefit from mutualism can be isolated as 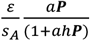; above carrying capacity (*K_A_* = *r_A_/s_A_*) that the animal population achieves by foraging on plant rewards. Benefits to the animal population saturate with increasing ***P*** due to time constraints from handling food rewards (*h*). This sets an upper bound (horizontal asymptote) on animal density at 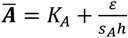. When facultative, the animal nullcline intersects the y-axis in the positive quadrant at animal carrying capacity 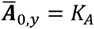. All else being equal, decreasing *r_A_* pushes the visible part of the animal nullcline down so that when the animal population becomes obligate, its nullcline intersects the y-axis at zero or negative values (not shown). Instead, for obligate animals the x-intercept becomes visible in the positive quadrant at

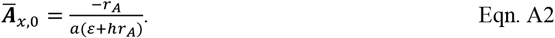

Plant dynamics are specific to each mutualism. The equation for the nullcline of animal-pollinated plants (Fig. 1, green) is:

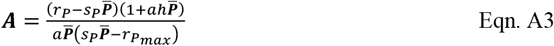

where *r_P_* = *b_P_ fg-d_P_* is the intrinsic growth rate of the plant population, ***P***, and *r_P_max__* = *r_P_* + *b_P_φg* is its maximum per-capita growth rate in the presence of pollinators. For feasible coexistence, *r_P_max__* > 0. The plant nullcline is bounded between vertical asymptotes at ***P*** = 0 and ***P*** = *r_P_max__/s_P_*, where ***P*** = *r_P_max__/s_P_* represents the plant’s maximum population density. When plants are obligate (*r_P_* ≤ 0, Fig. 1A-B), the nullcline is a U-shape. All else being equal, increasing *r_P_* pushes the minimum point of the U down towards the x-axis, so that when the plant population becomes facultative, it intersects, flipping into a cubic shape. Therefore, when plants are facultative (*r_P_* > 0, Fig. 1C-G), the nullcline is concave up at high density but concave down at low density, though this inflection may not be visible in the positive quadrant. The x-intercept is the plant carrying capacity 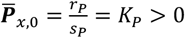. Inside the U or to the left of the plant nullcline, plant density increases; to the right or under, plant density decreases due to strong negative density-dependence.

The equation for the nullcline of animal-dispersed plants that benefit through reduced negative density-dependence (Fig. 2, green) is:

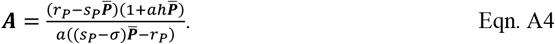

In the positive quadrant, the nullcline is increasing, concave up and saturates to a maximum plant density of ***P*** = *r_P_*/(*s_P_* – *σ*) when *s_P_* > *σ*. This maximum disappears to infinity when *s_P_* = *σ*, that is, when dispersers can completely remove sources of negative density-dependence. Thus, based on the nullcline alone, benefits to plants would increase indefinitely with increasing animal density. However, animal visitation rate saturates due to handling time (encoded in the Holling type II functional response), bounding the benefits to plants by intersecting the plant nullcline at high density. Because we consider only facultative plants for this mutualism, the nullcline always has an x-intercept at plant carrying capacity 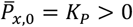 but no y-intercept in the positive quadrant.

The equation for the nullcline of animal-dispersed plants that benefit through germination (Fig. 3, green) is:

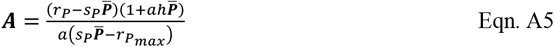

where *r_P_max__* = *r_P_* + *b_P_fγ* is the maximum per-capita growth rate of the plant population in the presence of dispersers. For feasible coexistence, *r_P_max__* > 0. In the positive quadrant, the nullcline is an increasing, concave up curve, bounded by a vertical asymptote at ***P*** = *r_P_max__/s_P_*, the maximum plant density in the presence of mutualism. The nullcline intersects the x-axis at plant carrying capacity 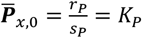 which is visible in the positive quadrant when plants are facultative (*r_P_* > 0). It intersects the y-axis at

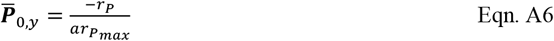

which is visible in positive quadrant when plants are obligate (*r_P_* ≤ 0).

### Note on Functional Forms

Because reproductive services provided by animals are a function of their consumption rate on plant rewards (*C_R_*, Eqn. 2), only one functional form per model needs to be specified. Pollination services 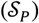 are assumed to be equal to animal’s total consumption rate on plants, while seed dispersal services 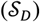 are assumed to be equal to the per-plant consumption rate. Reproductive services are assumed to saturate according to 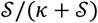, where *κ* is saturation coefficient for benefit from animal visitation rate with units [*A t*^-1^] or [*A P*^-1^ *t*^-1^] for pollination or seed dispersal mutualisms, respectively. For convenience, we set *k* = 1 in the main text.

